# Transcutaneous auricular vagus nerve stimulation during short-term motor practice drives cortical plasticity without behavioral improvement

**DOI:** 10.1101/2025.08.08.668810

**Authors:** Kento Nakagawa, Rieko Osu

**Author notes:** Corresponding author email address Kento Nakagawa Rieko Osu.

## Abstract

Transcutaneous auricular vagus nerve stimulation (taVNS) is emerging as a promising non-invasive neuromodulation technique to augment neurorehabilitation, yet its mechanisms in humans remain poorly understood. Animal studies suggest that VNS delivered during motor skill practice drives task-specific plasticity in the primary motor cortex (M1), but direct evidence in humans has been lacking. Here, we provide the first demonstration that taVNS paired with motor skill practice selectively enhances cortical plasticity without boosting motor performance beyond practice alone during short-term training. Thirty-one healthy right-handed adults practiced a novel implicit motor task, rotating two balls with the non-dominant hand for 15 minutes. Participants were randomized to receive taVNS to the left tragus or sham stimulation during practice. Motor performance, M1 hand representation (TMS mapping), and spinal excitability (F-wave) were assessed pre- and post-practice, while pupil diameter was continuously monitored as an index of noradrenergic activity. Motor performance improved similarly in both groups, whereas cortical map expansion was significantly greater in the taVNS group than in the sham group. F-wave amplitude increased only in the sham group, suggesting that taVNS-driven plasticity was restricted to cortical circuits. Moreover, taVNS uniquely elicited pupil dilation during practice, consistent with noradrenergic system engagement. These findings reveal that taVNS can promote task-specific cortical reorganization in humans independent of immediate behavioral improvement. By linking taVNS-induced plasticity to noradrenergic modulation and dissociating cortical from spinal effects, this study provides novel mechanistic insight into how taVNS may lay the neural groundwork for enhanced motor recovery, with critical implications for neurorehabilitation.

**Significance Statement:** Motor recovery after stroke depends on the brain’s ability to reorganize motor circuits. Vagus nerve stimulation (VNS) has been proposed as a powerful approach to enhance such plasticity, but the underlying mechanisms in humans remain unclear. Here we show that non-invasive transcutaneous auricular VNS (taVNS), when paired with motor practice, selectively enhances reorganization of the primary motor cortex without additional behavioral improvement. Importantly, these effects were observed after only 15 minutes of practice, demonstrating that taVNS can induce rapid plastic changes. This provides the first evidence in humans that taVNS promotes task-specific cortical plasticity, underscoring its promise as a neuromodulatory tool for neurorehabilitation.

## Introduction

Enhancing neuroplasticity of the central nervous system (CNS) is a key strategy for promoting neurorehabilitation and motor skill acquisition. Vagal nerve stimulation (VNS) has emerged as a promising tool to enhance motor recovery, as shown in both animal models and clinical trials in stroke patients (de Melo et al., 2023; Gerges et al., 2024; Korupolu et al., 2024).Transcutaneous auricular VNS (taVNS) provides a non-invasive, safe, and accessible alternative to invasive VNS and has therefore attracted growing therapeutic interest. Recent meta-analyses have demonstrated its safety taVNS (Kim et al., 2022) and suggested superior therapeutic effects compared to other neuromodulation techniques in stroke patients (Ahmed et al., 2023). However, the underlying therapeutic mechanisms of taVNS remain unclear.

In able-bodied individuals, taVNS has been reported to improve cognitive function (Sun et al., 2021), but no study has yet demonstrated improvement in motor skills with taVNS in able-bodied individuals. Animal studies have shown that invasive VNS delivered during dexterous motor training induces plastic changes in the primary motor cortex (M1) representation in a task-dependent manner (Porter et al., 2012; Morrison et al., 2019; Brougher et al., 2021). This suggests that pairing taVNS with motor skill training may facilitate plastic changes in motor-related neural circuits innervating task-relevant muscles. In human, studies examining effects of taVNS in the absence of a motor task on M1 excitability or inhibition have yielded inconsistent results (Mertens et al., 2022; Gerges et al., 2025; Yun et al., 2025). Notably, no study has yet assessed plastic changes in M1 motor map representation following taVNS. As M1 neurons project directly to spinal motoneurons, cortical activation or even subthreshold plastic changes in M1 may also influence spinal excitability. Indeed, non-invasive stimulation over M1 has been shown to modulate not only cortical excitability but also spinal circuits (Kondo et al., 2013; Klomjai et al., 2022). As voluntary motor execution requires regulation of spinal motoneuron excitability, taVNS delivered during motor learning could potentially affect both cortical and spinal motoneuron. However, the impact of taVNS on spinal motoneuron excitability remains unexplored.

taVNS has also been shown to modulate autonomic and arousal systems. Studies have suggested that taVNS in rest activates the locus coeruleus-noradrenaline (LC-NA) system, as reflected by pupil dilation—a reliable biomarker of LC-NA activity (Sharon et al., 2021; Lloyd et al., 2023; Pervaz et al., 2025).Yet it remains unclear whether taVNS paired with motor execution engages the same neural circuits as taVNS at rest. Monitoring pupil dynamics during taVNS paired with motor learning may therefore provide insights into underlying neuromodulatory mechanisms.

Previous work in both animals and humans has highlighted the potential importance of stimulation timing. In several studies, phasic VNS was delivered concurrently with discrete movements such as reaching, grasping, and lever pressing (Porter et al., 2012; Khodaparast et al., 2014; Dawson et al., 2021). An animal study further suggested enhanced motor improvements and selective modulation of M1 neurons when VNS delivers immediately after successful movements (Bowles et al., 2022). Meanwhile, other rehabilitation studies reported beneficial effects of taVNS regardless of precise timing, for example when continuous taVNS at rest was combined with subsequent therapy (Wu et al., 2020; Li et al., 2022). Thus, the importance of stimulation timing remains unresolved. Moreover, most previous studies have employed discrete tasks, leaving it unknown whether taVNS can modulate learning of periodic motor skills. To address this gap, we employed a periodic two-ball rotation task as the experimental model. This task recures sequential and coordinated finger movements, and has been widely used in motor learning paradigm (Uehara et al., 2011; Hamada et al., 2023). Although the objective of learning this skill is to rotate the balls faster, each rotation can be regarded as a successful movement, even if performed slowly. Thus, continuous taVNS during practice of two-ball rotation can be regarded as stimulation delivered concurrently with ongoing successful movements, analogous to phase VNS paired with discrete successes in prior studies.

In this study, we investigated the effects of taVNS paired with periodic motor skill practice on behavioral performance and plasticity in motor-related neural circuits. We hypothesized that taVNS during motor practice would facilitate motor learning, expand M1 representations of task-relevant muscles, and increase spinal motoneuron excitability, potentially through LC-NA system engagement.

## Methods

### Participants

Thirty-one healthy right-handed adults participated in this experiment. Participants were pseudo-randomly assigned to either the taVNS group (n=16; 23 ± 4 years; 3 females and 13 males] or the sham group (n=15; 22 ± 5 years; 4 females and 11 males). Handedness was confirmed using the Edinburgh Handedness Inventory (Oldfield, 1971), with laterality quotient ranging from 68.4 to 100 (mean ± SD: 93.7 ± 10.2). None of the participants had prior experience with the two-ball rotation task. This study was approved by the Human Research Ethics Committee of Waseda University (approval number: 2020-93), and written informed consent was obtained from all participants.

### Overall design and procedure

This study employed a randomized, single-blind, sham-controlled design. Participants in both groups were told that electrical stimulation would be delivered to the ear electrodes, but that the current might be too weak to perceive. Thus, all participants believed they were receiving real stimulation, regardless of group assignment. Since the stimulator did not include a sham mode and the experimenter manually adjusted the stimulation intensity, the experimenter was not blinded.

Figure 1 summarizes the experimental protocol. The procedure was identical for both group except for the presence (taVNS group) or absence (sham group) of electrical stimulation on ear electrode during motor practice. Participants performed a two-ball rotation task with the left hand, consisting of 30 practice trials (10 trials × 3 sets). During practice, taVNS or sham stimulation was applied to the tragus. Behavioral performance and neurophysiological assessments were conducted immediately before and after practice sessions (Pre- and Post-tests) without taVNS electrodes attached. Each experimental session lasted approximately three hours.

**Figure 1.**
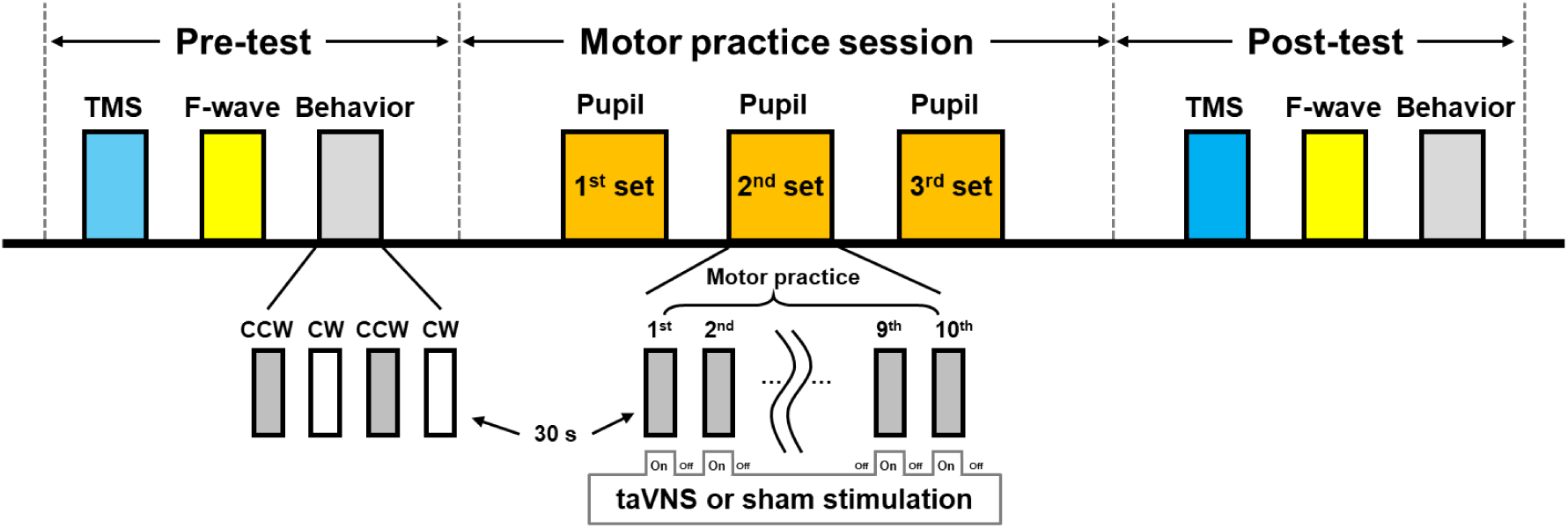
Experimental protocol. In the Pre- and Post-test, TMS mapping, F-wave measurement, and behavioral assessments were conducted. Behavioral tests consisted of 30-s trials of clockwise (CW) and counterclockwise (CCW) ball rotation. During the practice session, participants performed 30 trials of 30-s CW rotation with 30-s rest intervals. taVNS or sham stimulation was applied during practice.

### taVNS

During motor practice, participants received either taVNS or sham stimulation (no current). To target the afferent auricular branch of the vagus nerve, a stimulation electrode (RELIfit-Tragus, Soterix Medical Inc.) was secured by hooking the device over the left ear, with the tip of the electrode positioned on the tragus (Figure 2). The position was adjusted to maximize perceptual intensity and fixed with medical tape. The skin around the tragus was cleaned with alcohol wipes before placement. The tragus was chosen because it has been suggested as a suitable site for vagal modulation (Butt et al., 2020).

Stimulation was delivered using a dedicated device (0125tA, Soterix Medical Inc.) with parameters based on previous studies (Wu et al., 2020; Chang et al., 2021; Li et al., 2022): 30 Hz frequency, 300 μs pulse width, and a total duration of 900 s (30 blocks of 30-s on and 30-s off). Current intensity was set at the individual pain threshold or 0.5 mA below, because the device allowed only 0.5 mA increments. If stimulation at the pain threshold was reported as “intolerable,” intensity was reduced by 0.5 mA, which still exceeded the perception threshold in all participants. The mean stimulation intensity was 2.37 ± 1.41 mA. For sham stimulation, some previous studies applied current to the ear lobe (Wu et al., 2020; van Midden et al., 2023), where the vagal innervation is absent. However, as ear lobe stimulation may still influence motor-related brain functions (van Midden et al., 2023), we used a no current sham condition with the electrode placed at the same tragus position, consistent with other studies (Dawson et al., 2021; Li et al., 2022). No participants reported adverse effects such as headaches.

**Figure 2.**
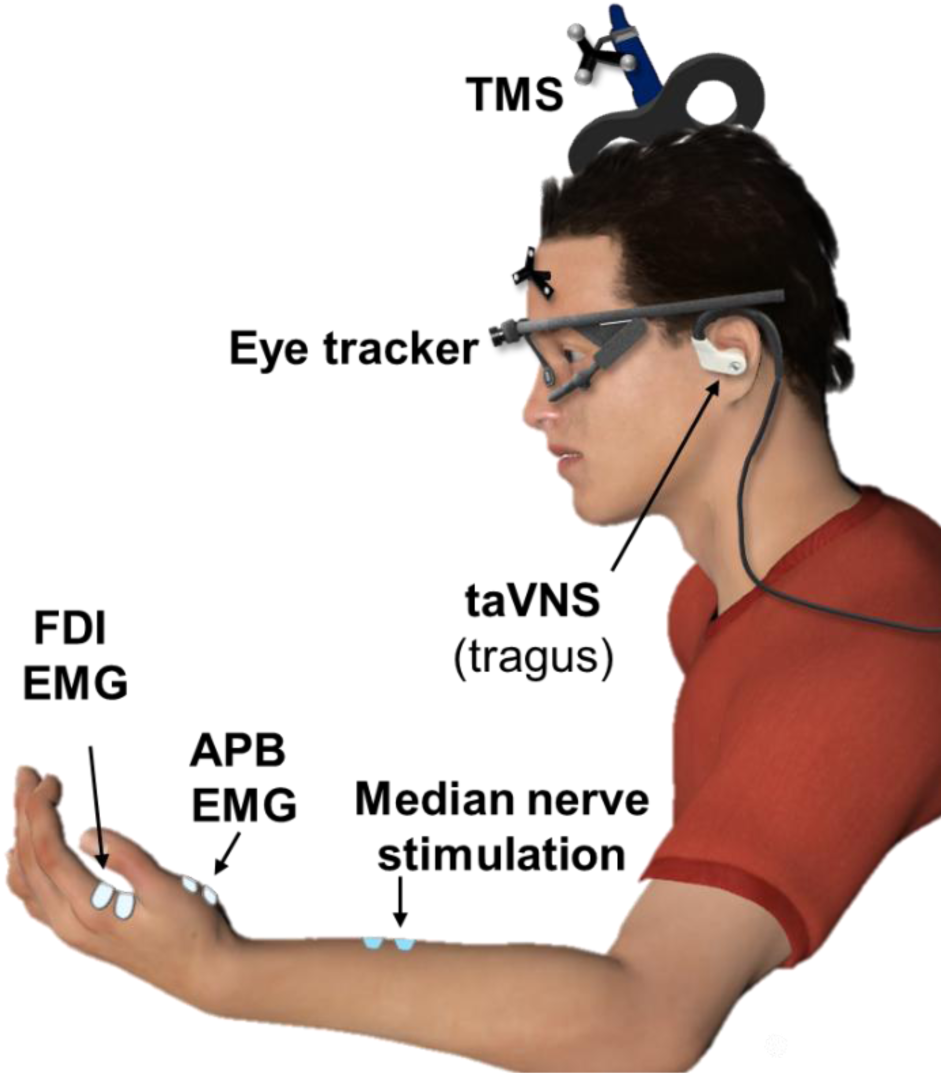
Experimental setup. The human figure was computer-generated using Poser 12 software and does not depict any real person. The equipment shown was created by the authors based on the actual apparatus used in the experiment

### Motor task and practice

Participants practiced a periodic two-ball rotation task with their left hand (Figure 3A). Seated at a desk, they rotated two golf balls (33 mm in diameter, 45g each; D1 Plus, Homa, Japan) placed in the supine left palm as quickly as possible in a clockwise direction. The balls were colored yellow and white to facilitate tracking during later video analysis. Participants were instructed to maintain their gaze so that both the left hand and the monitor displaying task instructions were within view. If a ball was dropped, participants were instructed to pick it up immediately and continue the task.

During the Pre- and Post-test, participants performed two 30-s trials of the practiced clockwise (CW) rotation and two 30-s trials of the unpracticed counterclockwise (CCW) rotation, each performed as fast as possible. Regardless of test or practice, every 30-s motor task was preceded and followed by a 30-s rest period without practice or stimulation. One practice set consisted of 10 alternating trials of 30-s practice and 30-s rest. Participants completed three sets in total, with breaks between sets determined individually (typically 1–2 min). Task instructions, trial timing, and elapsed time were displayed on the monitor to ensure standardized practice across participants.

For behavioral assessment, performance was recorded using a webcam (D600, EMEET, China) mounted on a tripod (Q111H, Say Good) above the participant’s hand, at a frame rate of 60 fps. Video data were analyzed with DeepLabCut, an open source markerless pose estimation toolbox with deep neural networks (Mathis et al., 2018). The y-axis displacement of one tracked ball was used to compute two parameters in MATLAB (Mathworks, USA): (1) the number of rotations, determined by peak cycle counts, and 2) rotation instability quantified as the standard deviation of cycle length.

### TMS measurement

Electromyography (EMG) signals were recorded from the left abductor pollicis brevis (APB) and first dorsal interosseous muscle (FDI) using bipolar Ag/AgCl surface electrodes (Vitrode F-150S, 18 × 36 mm, Nihon Kohden, Tokyo, Japan) (Figure 2). EMG signals were amplified (MEB-6108 amplifier, Nihon Kohden, Tokyo, Japan), band-pass filtered (5–1500 Hz), and digitized at 4000 Hz with an A/D converter (Powerlab, AD Instrument, Sydney, Australia). Single-pulse monophasic TMS was delivered with a Magstim 200 (Magstim, Whitland, UK) through a 70-mm figure-eight coil. Coil position, orientation, and tilt relative to the head were continuously monitored with a navigation system (Brainsight version 2.2.8, Rogue Research, Montreal, Canada). Participants were seated comfortably and instructed to remain relaxed throughout the measurements.

The coil was positioned over the site that elicited the largest motor evoked potential (MEP) in the APB, designed as the hotspot. The coil orientation and tilt at the hotspot were recorded to ensure reproducibility. Resting motor threshold (RMT) was defined as the lowest TMS intensity that evoked MEPs > 0.05 mV in at least 5 of 10 trials in the APB.

Motor maps were then obtained using TMS at120% RMT within an 11×11 cm grid (1 cm spacing) centered on the hotspot (Figure 4A). Coil orientation was held constant relative to the hotspot, while tilt was adjusted to maintain tangential contact with the scalp. Three TMSs were delivered at each grid point every 3-4 s, guided by the navigation system. Mapping began at the hotspot and expanded outward. The next stimulating site was randomly chosen from one of the four neighbors of active points (mean APB MEP amplitude > 0.05 mV) that had not yet been tested. Sites with mean APB MEP < 0.05 mV were defined as inactive. Mapping continued until the entire map was surrounded by inactive sites.

MEP amplitude was quantified as the peak-to-peak value within a 20–50 ms windows after TMS (van de Ruit et al., 2015). Motor map characteristics were evaluated in two ways: (1) map size, defined as the number of active points, which reflects the spatial extent of the representation (Nakagawa et al., 2020; Maeo et al., 2021), and (2) map volume, defined as the sum of the mean MEP amplitudes across all active points, providing information on the overall excitability of the cortical representation (Rossini et al., 2015; Maeo et al., 2021).

### F-wave measurement

F-waves were elicited by electrical stimulation of the median nerve approximately 8 cm proximal to the left wrist, using 3.2-cm circular electrodes (cathode distal, anode proximal) connected to a constant-current stimulator (DS7A, Digitimer, UK) (Figure 2). The inter-electrode distance was approximately 5 cm. Prior to electrodes attachment, the optimal stimulation site was identified with a probe electrode as the location producing clear M-waves in both APB and FDI muscles. Participants were seated comfortably with the left hand supinated on a desk.

For each test (Pre- and Post-test), twenty stimuli (200 μs pulse width) were delivered at 0.5 Hz at 120% of the intensity required to evoke the maximum M-wave amplitude (M-max) in the APB. One participant in the taVNS group did not undergo F-wave measurement due to pain intolerance.

As F-wave latency varies across and within participants, the analysis window for peak-to-peak amplitudes was adjusted individually for each trial by visual inspection. Spinal excitability was assessed using two parameters: (1) F-wave persistence, defined as the percentage of detectable F-waves out of 20 stimuli (detection threshold: 0.03 mV) (Wupuer et al., 2013), and (2) F-wave amplitude, defined as the mean peak-to-peak amplitude of detectable F-waves normalized to M-max (F/M ratio). If no F-wave was detected across the 20 trials, amplitude data for that participant was excluded from statistical analysis.

### Statistical analysis of behavior, MEP, and F-wave

Data normality was assessed using the Shapiro–Wilk test. When normality was confirmed, two-way repeated measures ANOVAs (group [taVNS, sham] × time [Pre, Post]) were performed. When a significant interaction was found, post-hoc tests were performed: unpaired t-tests for between-group comparisons (taVNS vs. sham) and paired t-tests for within-group comparisons (Pre vs. Post) For these datasets with four comparisons, the significant level was adjusted to p < 0.0125 using Bonferroni correction. When normality was violated, non-parametric tests were applied (Mann– Whitney U-test for between-group comparisons and Wilcoxon signed-rank test for within-group comparisons).

Additionally, the magnitude of change (Δ) from Pre-to Post-test was calculated for each parameter. Between-group comparisons of Δ values were performed using unpaired t-tests (for normally distributed data) or Mann–Whitney U-tests (for non-normal data). Within-group Δ values were compared against zero using one-sample t-tests (parametric) or Wilcoxon signed-rank tests (non-parametric). For analyses of Δ values, significance was set at p < 0.05.

### Measurements and analysis of pupillary response

During the practice session (task and rest periods), pupil diameter was recorded at 120 Hz from the left eye using an eye tracker (Pupil Core, Pupil Labs, Germany) (Figure 2). The tracker was mounted immediately before the first practice set and removed after the final set.

Pupil data were preprocessed in MATLAB R2021a following the guidelines of Kret and Sjak-Shie (2019). Only samples with confidence value ≥ 0.7 were retained. Values outside the physiologically plausible range of 1.5–9 mm were discarded. Blink-related artifacts were identified using the first derivative of the pupil signal (dilation speed). Sample exhibiting the median + 3 median absolute deviations (MAD) were classified as artifact, and an additional 100 ms (three samples before and after) was removed to capture the full blink. Continuous gaps > 75 ms were expanded by 50 ms on each side. Further noise artifacts were detected by comparing the raw signal with a moving-average smoothed version; deviation > 3 MAD were excluded. Missing data were linearly interpolated only for gaps ≤ 0.5 s. The resulting signal was resampled to 30 Hz and smoothed with a zero-phase low-pass Butterworth filter (4 Hz cutoff).

Data were segmented into task and rest periods. For each period, pupil diameter was baseline corrected to the mean of the 0.2 s preceding the task/rest switch (stimulation on/off in taVNS group) (Wienke et al., 2023) and expressed as percentage change. Values were averaged across the 30 task or rest periods for each participant.

Time series differences in pupil diameter were examined using Statistical Parametric Mapping for one-dimensional data (SPM1D, www.spm1d.org) implemented in MATLAB (Meissner et al., 2024). One-sample t-tests against baseline (0) were performed within each condition. Paired t-tests compared task versus rest within each group, and unpaired t-tests compared taVNS and sham group under the same condition. At each time point, t- and p-values were computed, with significance set at α = 0.05 (two-tailed). To ensure physiological relevance, only clusters > 3 consecutive time points (> 0.1 s at 30 Hz; Wierda et al., 2012) exceeding the critical threshold were considered significant. Analysis were restricted to 0.2 – 25.0 s of 30-s trial, excluding the first 0.2 s (latency of pupil response) (Mathôt, 2018) and the final 5 s (anticipatory effects from focusing on the timer or task completion). In addition, the velocity of the change of pupil size was compared between the groups to assess potential differences in response dynamics. Two participants (taVNS group: N = 1, sham group: N = 1) were excluded due to eye tracker malfunction.

## Results

### Motor performance (Fig. 3)

Figure 3B shows representative displacement traces of the two balls along the y-axis during clockwise (CW) rotation in the Pre- and Post-tests. Compared with the Pre-test, rotation velocity and stability increased in the Post-test.

For the number of rotations, Shapiro–Wilk tests confirmed normality in all datasets. Two-way ANOVAs revealed a significant main effect of time (CCW: F(1, 29) = 49.397, p < 0.001; CW: F(1, 29) = 127.179, p < 0.001), but no significant main effect of group or group × time interaction (Fig. 3C).

For the rotation instability, non-normality was detected in several datasets (e.g., taVNS_Post in CCW, sham_Pre in CCW, taVNS_Pre in CW. Subsequent non-parametric analyses showed that instability in CW rotations significantly decreased from Pre- to Post-test in both the taVNS (p < .001) and sham groups (p < .001). No significant difference between taVNS and sham groups at either Pre- and Post-test. For CCW rotations, instability significantly decreased from Pre- to Post-test in the taVNS group (p = 0.008) but not in the sham group (Fig. 3D).

**Figure 3.**
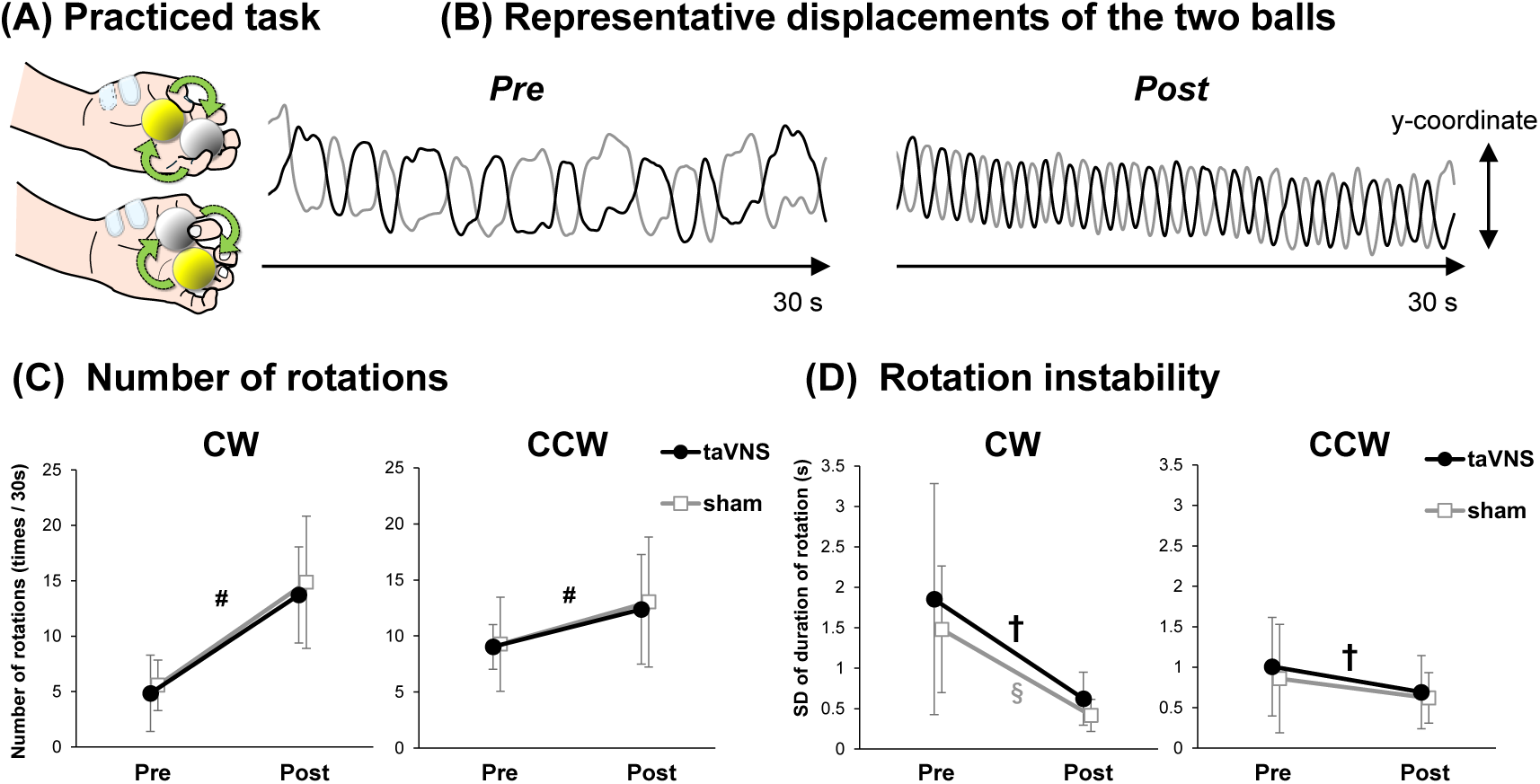
Motor task and behavioral results. (A) Practiced motor task: Participants rotated two balls clockwise (CW) in the left palm. In Pre- and Post-tests, non-practiced counterclockwise (CCW) rotation was also assessed. (B) Representative y-axis displacement of the two balls (black: white ball; gray: yellow ball) digitized with DeepLabCut during 30-s CW rotation in Pre- and Post-tests. (C) Group data for the number of rotations (Pre vs. Post) in each group (taVNS, sham). Both groups improved in CW and CCW, with no significant group differences. (D) Group data for rotation instability (SD of cycle duration). In CW, instability decreased in both groups; in CCW, only the taVNS group showed a decrease. Error bars = ±1 SD. # Significant main effect of time; † Significant Pre-Post difference in taVNS group; § Significant Pre-Post difference in sham group.

### Motor representation by TMS mapping

#### Map size (Fig. 4C)

For the APB muscle, Shapiro–Wilk test indicated non-normality in the taVNS_Pre and taVNS_Post datasets. No significant differences were detected in pairwise comparisons. For the Δ map size, normality was confirmed in both groups, and unpaired t-tests revealed that Δ map size was significantly greater in the taVNS group than in the sham group (p =.046).

For the FDI muscle, non-normality was detected in the taVNS_Post datasets, and no significant differences were found in pairwise comparisons. For the Δ map size, normality was confirmed in both groups, but no significant group differences were observed.

#### Map volume (Fig. 4D)

For the APB muscle, non-normality was observed in the taVNS_Pre and taVNS_Post datasets, and no significant differences were found in pairwise comparisons. For the Δ map volume, normality was not confirmed in the taVNS group. Mann-Whitney U-test showed that Δ map volume was significantly greater in the taVNS group compared with the sham group (p =.024).

For the FDI muscle, non-normality was detected in the taVNS_Pre and taVNS_Post datasets, with no significant pairwise differences. For the Δ map volume, normality was not confirmed in the taVNS group. Mann-Whitney U-test indicated that Δ map volume was significantly greater in the taVNS group than in the sham group (p =.017). In addition, Wilcoxon signed-rank test showed that Δ map volume in the sham group was significantly less than zero (p = .017).

**Figure 4.**
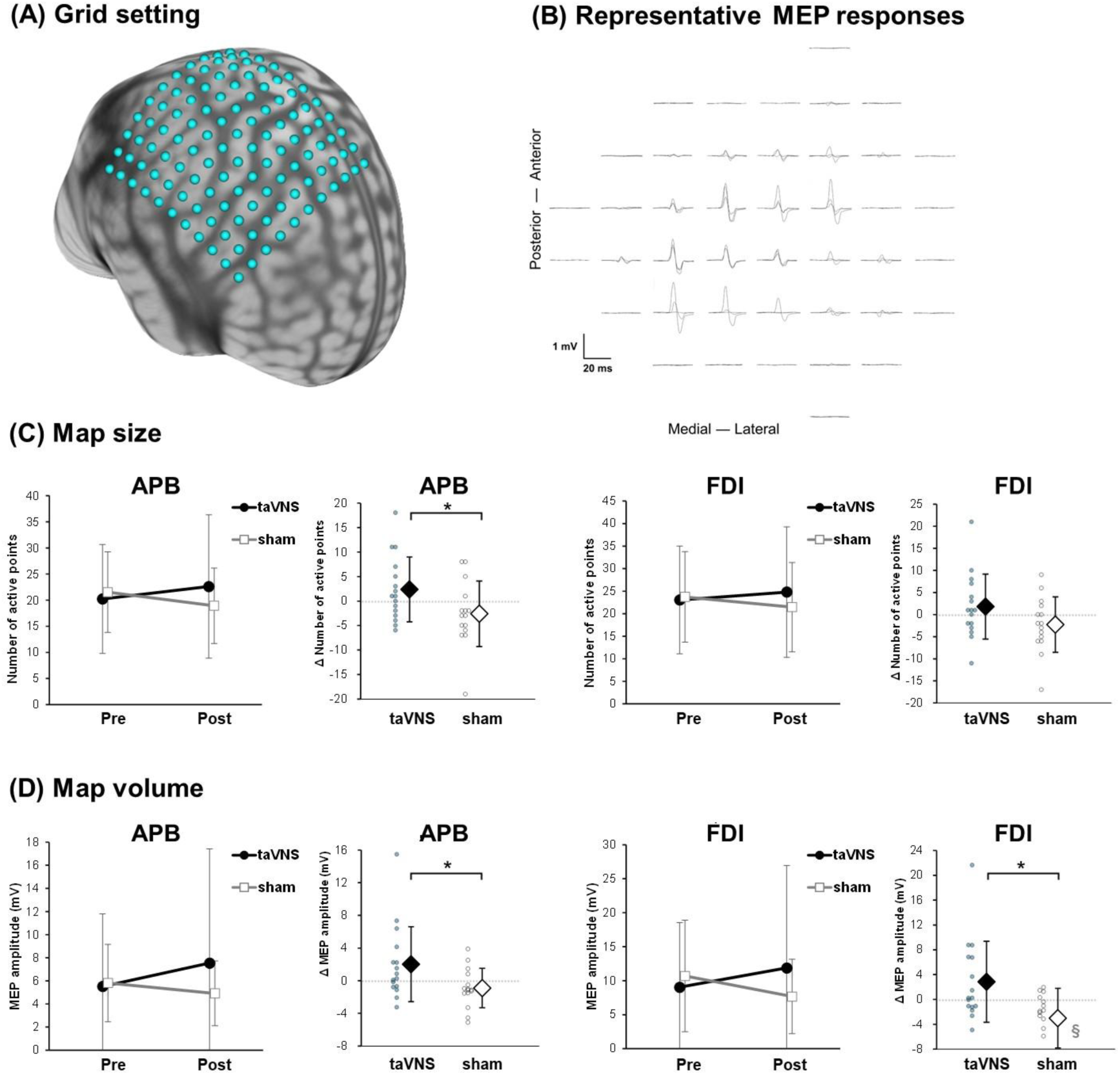
TMS motor mapping and results. (A) 11 × 11 grid centered on APB hotspot (B) Representative APB MEPs from one participant in the Pre-test (3 traces per grid point). (C) Group data for the map size (number of active points). (C) Group data for the map volume (sum of mean MEP amplitudes at active points). Error bars = ±1 SD. * p < 0.05, taVNS vs. sham; § p < 0.05, Δ vs. zero.

### Spinal excitability by F-wave measurements

#### F-wave persistence (Fig. 5B)

For the APB muscle, Shapiro–Wilk test indicated non-normality in several datasets (taVNS_Post, sham_Pre, and sham_Post). No significant differences were found in pairwise comparisons. For the Δ persistence, normality was confirmed in both groups, and unpaired t-tests showed no significant group differences.

For the FDI muscle, non-normality was detected in the sham_Post dataset, but no significant differences were observed in pairwise comparisons. Similarly, Δ persistence values showed no significant group differences.

#### F-wave amplitude (Fig. 5C)

For the APB muscle, all datasets met normality assumptions. A two-way ANOVA revealed a significant group × time interaction (F(1, 27) = 6.49, p = .017), with no main effects of group or time. However, follow-up paired t-tests did not identify significant differences among the four individual conditions. For Δ amplitude, unpaired t-tests showed that values were significantly greater in sham group compared with the taVNS group (p = .016). Additionally, the sham group exhibited a significant increase in Δ amplitude compared with zero (p= .034).

For the FDI muscle, normality was confirmed in all datasets, and no significant differences were detected in either pairwise comparisons or Δ amplitude values.

**Figure 5.**
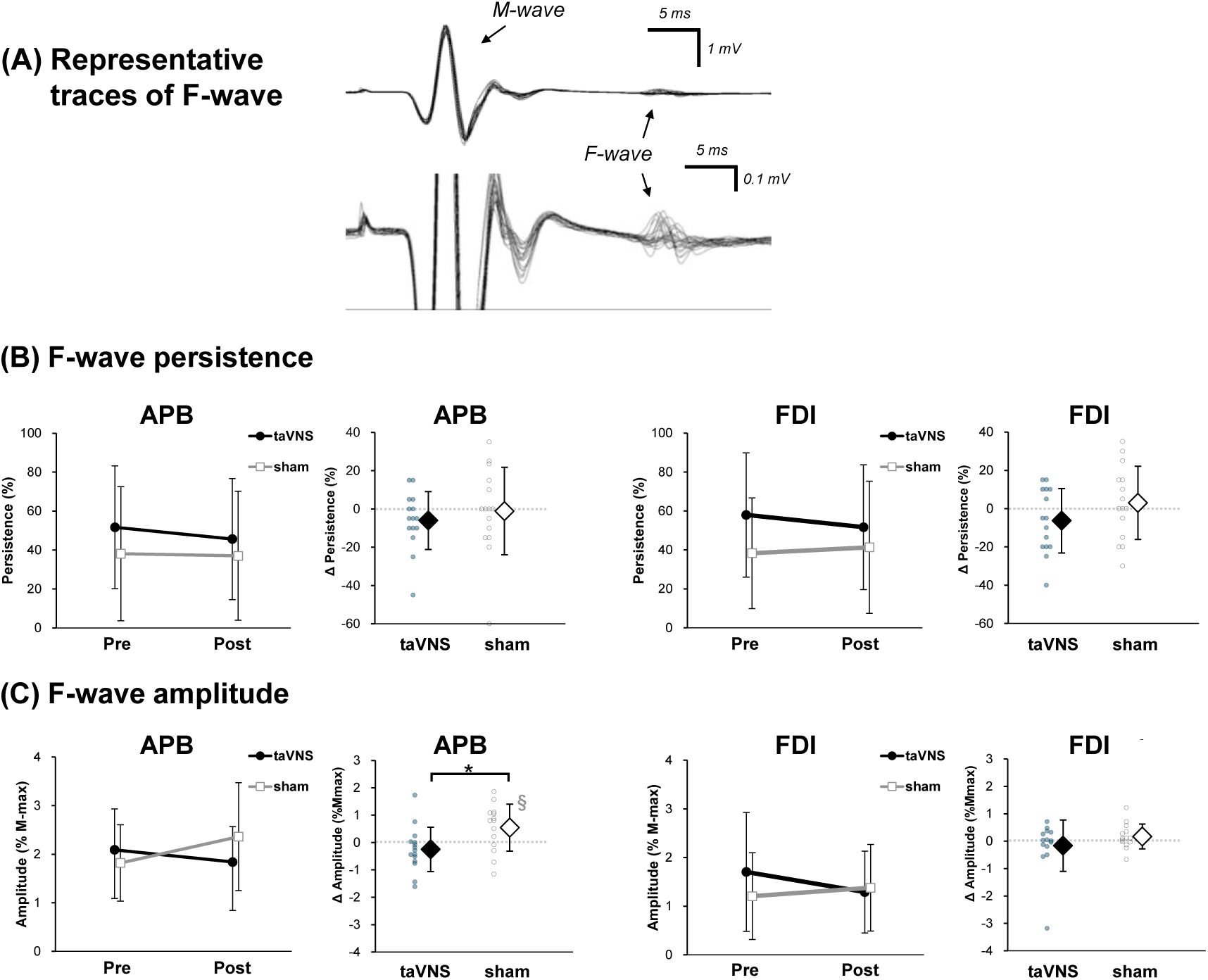
F-wave measurement and results. (A) Representative M-wave and F-wave traces from APB (20 trials, Pre-test). (B) Group data for F-wave persistence. (C) Group data for F-wave amplitude (F/M ratio). Error bars = ±1 SD. * p < 0.05, taVNS vs. sham; § p < 0.05, Δ vs. zero.

#### Pupil dilation (Fig. 6A)

Figure 6A shows normalized pupil diameter from the onset of task/rest periods (0 s) to 30 s. During the task period, the taVNS group exhibited a significant pupil dilation relative to baseline between 0.5–1.9 s (*p* < .05), whereas no significant change was observed in the sham group. During the rest period, both groups showed significant pupil constriction relative to baseline: 1.3–25.0 s for taVNS and 3.7–25.0 s for sham group (*p* < .05). Direct comparisons between task and rest revealed significant differences in the VNS group across 0.5–17.4 s, 17.8–23.4 s, and 24.3–24.5 s, and in the sham group across 3.4–9.3 s, 9.7–15.2 s, 16.0–19.3 s, and 19.4–25.0 s (all *p* < .05).

#### Velocity of pupil dilation (Fig. 6B)

Figure 6B illustrates the velocity of pupil size changes during the first 5 s after onset. During the task period, the taVNS group showed significantly faster pupil dilation than the sham group at 0.3–0.4 s after onset (*p* < .05).

**Figure 6.**
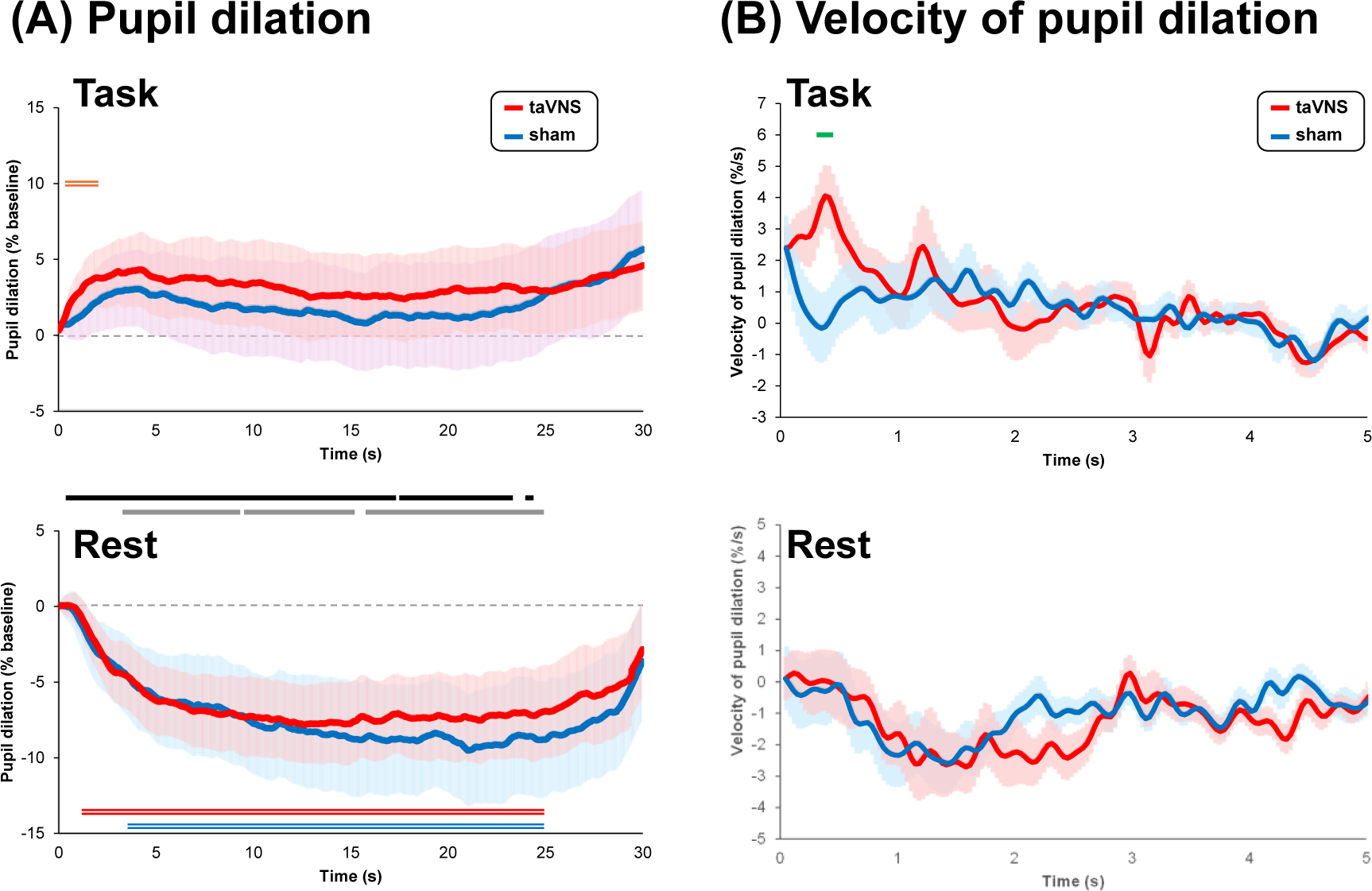
Pupil dilation during task and rest. (A) Grand-average pupil diameter (% change from baseline) during task (top) and rest (bottom). Solid lines: taVNS (red), sham (blue); shaded areas = ± SEM. Red double lines indicate time ranges significantly different from baseline in the taVNS group (p < 0.05); blue double lines indicate those in the sham group (p < 0.05). Black and gray horizontal bars denote periods where task–rest differences were significant within the taVNS and sham groups, respectively (p < 0.05). No significant differences were found between groups. (B) Grand-average of velocity of pupil dilation in the early phase (0–5s). Green bar indicates the period with a significant difference between groups during the task. No significant group differences during rest.

## Discussion

The objective of this study was to investigate whether short-term taVNS paired with periodic motor skill practice enhances motor learning and induces plasticity in motor-related neural circuits. Our results show that taVNS during two-ball rotation practice facilitated expansion of M1 representation of the finger muscles, whereas motor learning itself was not enhanced. Moreover, spinal motoneuron excitability increased in the sham group but not in the taVNS group. Finally, taVNS group exhibited stronger pupil dilation during practice, consistent with greater recruitment of the LC-NA system.

### Effects of taVNS on motor learning

Both groups significantly improved in two-ball rotation performance, but taVNS did not provide additional behavioral benefit. Thus, our hypothesis was not supported at the behavioral level. The most likely reason is the brevity of the intervention: only 15 min of practice with stimulation. In contrast, clinical studies that demonstrated functional gains in stroke patients typically involved intervention periods ranging from days to weeks (Gerges et al., 2024). Our findings suggest that the present protocol was sufficient to induce neural plasticity but insufficient to drive behavioral gains. Consistently, an animal study reported that VNS paired with motor training induced expansion of M1 maps without parallel performance gains (Hulsey et al., 2016), implying that cortical reorganization and behavioral improvements can occur on different timescales. Longer interventions may reveal behavioral benefits.

A second possibility is that the continuous ball-rotation task did not provide explicit signals of success, unlike paradigms where VNS was paired with discrete, rewarded movements (Bowles et al., 2022). Although each rotation could technically be considered as a success, participants may not have perceived trial-by-trial outcomes, reducing the engagement of reward-related neuromodulatory circuits. Motivation and task salience, typically high in rehabilitation or animal models, may have been relatively low here, attenuating the neuromodulatory influence of taVNS.

Interestingly, only the taVNS group improved stability in the non-practiced CCW condition. Given the greater M1 expansion observed in this group, this effect may reflect generalized learning through M1 reorganization, consistent with reports that non-invasive M1 stimulation promotes intracortical disinhibition and motor generalization (Dumel et al., 2018).

### Effects of taVNS on M1 plasticity

Our results clearly demonstrated greater expansion of M1 representations in the taVNS group than in the sham group, supporting our cortical-level hypothesis. This finding aligns with prior animal studies (Porter et al., 2012; Hulsey et al., 2016) and provides the first evidence in humans that taVNS can enhance motor map plasticity. Notably, the intervention duration was much shorter than in previous animal or clinical studies (days to weeks) (Porter et al., 2012; Hulsey et al., 2016; Gerges et al., 2024), highlighting the potency of taVNS to induce rapid cortical reorganization, which is highly relevant for clinical translation. In the sham group, most M1 parameters did not change significantly, except for a reduction in FDI map volume. This may reflect neural efficiency, a form of cortical plasticity where reduced activation supports improved skill execution (Callan and Naito, 2014). Therefore, taVNS may shift plasticity toward representational enlargement. Whether this expansion is muscle-specific or broader remains unclear, as only APB and FDI were examined.

### Effects of taVNS on spinal motoneuron excitability

To explore plasticity beyond the brain, measured F-waves as an index of spinal motoneuron excitability—a novel approach in taVNS or implanted VNS research. In the sham group, F-wave amplitude increased in APB, indicating use-dependent plasticity at the spinal level. Such spinal adaptations have been reported after training of cyclic or postural skills (Mazzocchio et al., 2006; Harel et al., 2015), where spinal contributions are likely substantial compared with the case of discrete movement. Thus, motor learning can drive both cortical and spinal plasticity.

In contrast, taVNS prevented these spinal changes. This suggest that taVNS restricts plasticity to supraspinal circuits, likely via selective engagement of vagal afferents that project to brainstem nuclei but not directly to the spinal cord. The modulation of cortical circuits may overshadow or cancel use-dependent spinal modulation. Consequently, even if behavioral improvements did not differ, the underlying neural substrates of learning were fundamentally different between groups.

### Effects of taVNS on pupil size

To our knowledge, this is the first study to examine pupil dynamics during motor practice with taVNS. Both groups showed task-related pupil dilation relative to rest, but only the taVNS group exhibited a rapid and significant dilation immediately after task onset, accompanied by faster dilation velocity than sham. Interestingly, pupil dilation also began several seconds before the switch from rest to task, likely reflecting predictive preparation, as the timing of stimulation onset was displayed on the monitor. Consequently, pupil size during the task period differed little from the immediate pre-task baseline, except for a brief interval of significant dilation observed only in the taVNS group. Pupil diameter and its rate of change are reliable markers of LC-NA system activity (Reimer et al., 2016), and thus these results indicate stronger noradrenergic engagement under taVNS. Because pupil size is also a proxy for cognitive states such as arousal, which are modulated by the LC-NA system during exercise and motor learning (Kuwamizu et al., 2022; Yokoi and Weiler, 2022), it is possible that taVNS participants were in a state of greater attentional focus on the task, which may in turn have contributed to the observed M1 plasticity.

### Possible mechanisms of M1 plastic changes by combination of motor practice and taVNS

Previous studies demonstrated that the two-ball rotation task engages M1, premotor, supplementary motor, cerebellar regions (Matsumura et al., 2004; Park et al., 2008). In parallel, taVNS activates the locus coeruleus (LC) and cerebellum even at rest (Frangos et al., 2015; Badran et al., 2018), and modulates cerebello-thalamo-M1 pathways (van Midden et al., 2024). Thus, combined activation of cerebellar circuits during practice and taVNS may have facilitated M1 reorganization via the root of LC-cerebellum-thalamus-M1. Although short-term practice was insufficient to improve behavior, longer interventions may strengthen cerebellar internal models and promote learning.

At the molecular level, VNS has been shown to enhance BDNF expression (Hays et al., 2013). Recent animal work indicates that taVNS similarly increases cortical BDNF via cholinergic receptor (Li et al., 2020), while human studies suggest cholinergic recruitment by taVNS (Horinouchi et al., 2024). The cholinergic system has been identified as a key mediator of both motor learning (Bowles et al., 2022) and plastic expansion of M1 representation (Hulsey et al., 2016). In addition, noradrenergic engagement, as supported by our pupil findings, is likely critical (Lloyd et al., 2023; Pervaz et al., 2025; Sharon et al., 2021). Both LC-NA and basal forebrain cholinergic systems receive projections from the nucleus of the solitary tract, the first relay of vagal afferents (Rodenkirch et al., 2022). Thus, taVNS probably co-activates these systems, leading to increased release of neuromodulators and BDNF, thereby facilitating M1 plasticity.

## Conclusion

In summary, short-term taVNS paired with motor practice did not enhance motor learning but selectively promoted reorganization of M1 map while preventing spinal plasticity. Pupil dynamics indicated heightened noradrenergic engagement, suggesting differential recruitment of neuromodulatory circuits between groups. Although it was insufficient to boost behavioral improvement in the short term, taVNS may provide a neural substrate for long-term functional recovery when combined with extended training. These findings identify taVNS as a selective modulator of cortical plasticity in humans and support its potential as a targeted adjunct to neurorehabilitation.

## Acknowledgments

This study was supported by Grant-in-Aid for Scientific Research (B) (JSPS KAKENHI Grant Number 22H03498) to K. Nakagawa, Grant-in-Aid for Scientific Research (A) (JSPS KAKENHI Grant Number 21H04425) to R. Osu, and Grant-in-Aid for Scientific Research (A) (JSPS KAKENHI Grant Number 23H00459) to R. Osu.

## References

Ahmed I, Mustafaoglu R, Rossi S, Cavdar FA, Agyenkwa SK, Pang MYC, Straudi S (2023) Non-invasive Brain Stimulation Techniques for the Improvement of Upper Limb Motor Function and Performance in Activities of Daily Living After Stroke: A Systematic Review and Network Meta-analysis. Arch Phys Med Rehabil 104:1683–1697.

Badran BW, Dowdle LT, Mithoefer OJ, LaBate NT, Coatsworth J, Brown JC, DeVries WH, Austelle CW, McTeague LM, George MS (2018) Neurophysiologic effects of transcutaneous auricular vagus nerve stimulation (taVNS) via electrical stimulation of the tragus: A concurrent taVNS/fMRI study and review. Brain Stimul 11:492–500.

Bowles S, Hickman J, Peng X, Williamson WR, Huang R, Washington K, Donegan D, Welle CG (2022) Vagus nerve stimulation drives selective circuit modulation through cholinergic reinforcement. Neuron 110:2867–2885 e2867.

Brougher J, Sanchez CA, Aziz US, Gove KF, Thorn CA (2021) Vagus Nerve Stimulation Induced Motor Map Plasticity Does Not Require Cortical Dopamine. Front Neurosci 15:693140.

Butt MF, Albusoda A, Farmer AD, Aziz Q (2020) The anatomical basis for transcutaneous auricular vagus nerve stimulation. J Anat 236:588–611.

Callan DE, Naito E (2014) Neural processes distinguishing elite from expert and novice athletes. Cogn Behav Neurol 27:183–188.

Chang JL, Coggins AN, Saul M, Paget-Blanc A, Straka M, Wright J, Datta-Chaudhuri T, Zanos S, Volpe BT (2021) Transcutaneous Auricular Vagus Nerve Stimulation (tAVNS) Delivered During Upper Limb Interactive Robotic Training Demonstrates Novel Antagonist Control for Reaching Movements Following Stroke. Front Neurosci 15:767302.

Dawson J et al. (2021) Vagus nerve stimulation paired with rehabilitation for upper limb motor function after ischaemic stroke (VNS-REHAB): a randomised, blinded, pivotal, device trial. Lancet 397:1545–1553.

de Melo PS, Parente J, Rebello-Sanchez I, Marduy A, Gianlorenco AC, Kyung Kim C, Choi H, Song JJ, Fregni F (2023) Understanding the Neuroplastic Effects of Auricular Vagus Nerve Stimulation in Animal Models of Stroke: A Systematic Review and Meta-Analysis. Neurorehabil Neural Repair 37:564–576.

Dumel G, Bourassa M, Charlebois-Plante C, Desjardins M, Doyon J, Saint-Amour D, De Beaumont L (2018) Multisession anodal transcranial direct current stimulation induces motor cortex plasticity enhancement and motor learning generalization in an aging population. Clin Neurophysiol 129:494–502.

Frangos E, Ellrich J, Komisaruk BR (2015) Non-invasive Access to the Vagus Nerve Central Projections via Electrical Stimulation of the External Ear: fMRI Evidence in Humans. Brain Stimul 8:624–636.

Gerges ANH, Williams EER, Hillier S, Uy J, Hamilton T, Chamberlain S, Hordacre B (2024) Clinical application of transcutaneous auricular vagus nerve stimulation: a scoping review. Disabil Rehabil 46:5730–5760.

Gerges ANH, Graetz L, Hillier S, Uy J, Hamilton T, Opie G, Vallence AM, Braithwaite FA, Chamberlain S, Hordacre B (2025) Transcutaneous auricular vagus nerve stimulation modifies cortical excitability in middle-aged and older adults. Psychophysiology 62:e14584.

Hamada H, Wen W, Kawasaki T, Yamashita A, Asama H (2023) Characteristics of EEG power spectra involved in the proficiency of motor learning. Front Neurosci 17:1094658.

Harel NY, Martinez SA, Knezevic S, Asselin PK, Spungen AM (2015) Acute changes in soleus H-reflex facilitation and central motor conduction after targeted physical exercises. J Electromyogr Kinesiol 25:438–443.

Hays SA, Rennaker RL, Kilgard MP (2013) Chapter 11 - Targeting Plasticity with Vagus Nerve Stimulation to Treat Neurological Disease. In: Progress in Brain Research (Merzenich MM, Nahum M, Van Vleet TM, eds), pp 275–299: Elsevier.

Horinouchi T, Nezu T, Saita K, Date S, Kurumadani H, Maruyama H, Kirimoto H (2024) Transcutaneous auricular vagus nerve stimulation enhances short-latency afferent inhibition via central cholinergic system activation. Sci Rep 14:11224.

Hulsey DR, Hays SA, Khodaparast N, Ruiz A, Das P, Rennaker RL, 2nd, Kilgard MP (2016) Reorganization of Motor Cortex by Vagus Nerve Stimulation Requires Cholinergic Innervation. Brain Stimul 9:174–181.

Khodaparast N, Hays SA, Sloan AM, Fayyaz T, Hulsey DR, Rennaker RL, 2nd, Kilgard MP (2014) Vagus nerve stimulation delivered during motor rehabilitation improves recovery in a rat model of stroke. Neurorehabil Neural Repair 28:698–706.

Kim AY, Marduy A, de Melo PS, Gianlorenco AC, Kim CK, Choi H, Song JJ, Fregni F (2022) Safety of transcutaneous auricular vagus nerve stimulation (taVNS): a systematic review and meta-analysis. Sci Rep 12:22055.

Klomjai W, Giron A, Mounir El Mendili M, Aymard C, Pradat-Diehl P, Roche N, Katz R, Bayen E, Lackmy-Vallee A (2022) Anodal tDCS of contralesional hemisphere modulates ipsilateral control of spinal motor networks targeting the paretic arm post-stroke. Clin Neurophysiol 136:1–12.

Kondo T, Kakuda W, Yamada N, Shimizu M, Hagino H, Abo M (2013) Effect of low-frequency rTMS on motor neuron excitability after stroke. Acta Neurol Scand 127:26–30.

Korupolu R, Miller A, Park A, Yozbatiran N (2024) Neurorehabilitation with vagus nerve stimulation: a systematic review. Front Neurol 15:1390217.

Kret ME, Sjak-Shie EE (2019) Preprocessing pupil size data: Guidelines and code. Behav Res Methods 51:1336–1342.

Kuwamizu R, Yamazaki Y, Aoike N, Ochi G, Suwabe K, Soya H (2022) Pupil-linked arousal with very light exercise: pattern of pupil dilation during graded exercise. J Physiol Sci 72:23.

Li J, Zhang Q, Li S, Niu L, Ma J, Wen L, Zhang L, Li C (2020) alpha7nAchR mediates transcutaneous auricular vagus nerve stimulation-induced neuroprotection in a rat model of ischemic stroke by enhancing axonal plasticity. Neurosci Lett 730:135031.

Li JN, Xie CC, Li CQ, Zhang GF, Tang H, Jin CN, Ma JX, Wen L, Zhang KM, Niu LC (2022) Efficacy and safety of transcutaneous auricular vagus nerve stimulation combined with conventional rehabilitation training in acute stroke patients: a randomized controlled trial conducted for 1 year involving 60 patients. Neural Regen Res 17:1809–1813.

Lloyd B, Wurm F, de Kleijn R, Nieuwenhuis S (2023) Short-term transcutaneous vagus nerve stimulation increases pupil size but does not affect EEG alpha power: A replication of Sharon et al. (2021, Journal of Neuroscience). Brain Stimul 16:1001–1008.

Maeo S, Balshaw TG, Lanza MB, Hannah R, Folland JP (2021) Corticospinal excitability and motor representation after long-term resistance training. Eur J Neurosci 53:3416–3432.

Mathis A, Mamidanna P, Cury KM, Abe T, Murthy VN, Mathis MW, Bethge M (2018) DeepLabCut: markerless pose estimation of user-defined body parts with deep learning. Nat Neurosci 21:1281–1289.

Mathôt S (2018) Pupillometry: Psychology, Physiology, and Function. J Cogn 1:16.

Matsumura M, Sadato N, Kochiyama T, Nakamura S, Naito E, Matsunami K, Kawashima R, Fukuda H, Yonekura Y (2004) Role of the cerebellum in implicit motor skill learning: a PET study. Brain Res Bull 63:471–483.

Mazzocchio R, Kitago T, Liuzzi G, Wolpaw JR, Cohen LG (2006) Plastic changes in the human H-reflex pathway at rest following skillful cycling training. Clin Neurophysiol 117:1682–1691.

Meissner SN, Bachinger M, Kikkert S, Imhof J, Missura S, Carro Dominguez M, Wenderoth N (2024) Self-regulating arousal via pupil-based biofeedback. Nat Hum Behav 8:43–62.

Mertens A, Carrette S, Klooster D, Lescrauwaet E, Delbeke J, Wadman WJ, Carrette E, Raedt R, Boon P, Vonck K (2022) Investigating the Effect of Transcutaneous Auricular Vagus Nerve Stimulation on Cortical Excitability in Healthy Males. Neuromodulation 25:395–406.

Morrison RA, Hulsey DR, Adcock KS, Rennaker RL, 2nd, Kilgard MP, Hays SA (2019) Vagus nerve stimulation intensity influences motor cortex plasticity. Brain Stimul 12:256–262.

Nakagawa K, Takemi M, Nakanishi T, Sasaki A, Nakazawa K (2020) Cortical reorganization of lower-limb motor representations in an elite archery athlete with congenital amputation of both arms. Neuroimage Clin 25:102144.

Oldfield RC (1971) The assessment and analysis of handedness: the Edinburgh inventory. Neuropsychologia 9:97–113.

Park JW, Kwon YH, Lee MY, Bai D, Nam KS, Cho YW, Lee CH, Jang SH (2008) Brain activation pattern according to exercise complexity: a functional MRI study. NeuroRehabilitation 23:283–288.

Pervaz I, Thurn L, Vezzani C, Kaluza L, Kuhnel A, Kroemer NB (2025) Does transcutaneous auricular vagus nerve stimulation alter pupil dilation? A living Bayesian meta-analysis. Brain Stimul 18:148–157.

Porter BA, Khodaparast N, Fayyaz T, Cheung RJ, Ahmed SS, Vrana WA, Rennaker RL, 2nd, Kilgard MP (2012) Repeatedly pairing vagus nerve stimulation with a movement reorganizes primary motor cortex. Cereb Cortex 22:2365–2374.

Reimer J, McGinley MJ, Liu Y, Rodenkirch C, Wang Q, McCormick DA, Tolias AS (2016) Pupil fluctuations track rapid changes in adrenergic and cholinergic activity in cortex. Nat Commun 7:13289.

Rodenkirch C, Carmel JB, Wang Q (2022) Rapid Effects of Vagus Nerve Stimulation on Sensory Processing Through Activation of Neuromodulatory Systems. Front Neurosci 16:922424.

Rossini PM et al. (2015) Non-invasive electrical and magnetic stimulation of the brain, spinal cord, roots and peripheral nerves: Basic principles and procedures for routine clinical and research application. An updated report from an I.F.C.N. Committee. Clin Neurophysiol 126:1071–1107.

Sharon O, Fahoum F, Nir Y (2021) Transcutaneous Vagus Nerve Stimulation in Humans Induces Pupil Dilation and Attenuates Alpha Oscillations. J Neurosci 41:320–330.

Sun JB, Cheng C, Tian QQ, Yuan H, Yang XJ, Deng H, Guo XY, Cui YP, Zhang MK, Yin ZX, Wang C, Qin W (2021) Transcutaneous Auricular Vagus Nerve Stimulation Improves Spatial Working Memory in Healthy Young Adults. Front Neurosci 15:790793.

Uehara S, Nambu I, Tomatsu S, Lee J, Kakei S, Naito E (2011) Improving human plateaued motor skill with somatic stimulation. PLoS One 6:e25670.

van de Ruit M, Perenboom MJ, Grey MJ (2015) TMS brain mapping in less than two minutes. Brain Stimul 8:231–239.

van Midden VM, Pirtosek Z, Kojovic M (2024) The Effect of taVNS on the Cerebello-Thalamo-Cortical Pathway: a TMS Study. Cerebellum 23:1013–1019.

van Midden VM, Demsar J, Pirtosek Z, Kojovic M (2023) The effects of transcutaneous auricular vagal nerve stimulation on cortical GABAergic and cholinergic circuits: A transcranial magnetic stimulation study. Eur J Neurosci 57:2160–2173.

Wienke C, Grueschow M, Haghikia A, Zaehle T (2023) Phasic, Event-Related Transcutaneous Auricular Vagus Nerve Stimulation Modifies Behavioral, Pupillary, and Low-Frequency Oscillatory Power Responses. J Neurosci 43:6306–6319.

Wu D, Ma J, Zhang L, Wang S, Tan B, Jia G (2020) Effect and Safety of Transcutaneous Auricular Vagus Nerve Stimulation on Recovery of Upper Limb Motor Function in Subacute Ischemic Stroke Patients: A Randomized Pilot Study. Neural Plast 2020:8841752.

Wupuer S, Yamamoto T, Katayama Y, Motohiko H, Sekiguchi S, Matsumura Y, Kobayashi K, Obuchi T, Fukaya C (2013) F-wave suppression induced by suprathreshold high-frequency repetitive trascranial magnetic stimulation in poststroke patients with increased spasticity. Neuromodulation 16:206–211; discussion 211.

Yokoi A, Weiler J (2022) Pupil diameter tracked during motor adaptation in humans. J Neurophysiol 128:1224–1243.

Yun YJ, Myong Y, Oh BM, Song JJ, Kim CK, Seo HG (2025) Effects of Transcutaneous Auricular Vagus Nerve Stimulation on Cortical Excitability in Healthy Adults. Neuromodulation 28:115–122.

